# Potential influence of school-based lifestyle strategies among Australian Aboriginal and non-Aboriginal children: a cross-sectional comparison of adiposity and weight related behaviours between 2010 and 2015

**DOI:** 10.1101/518233

**Authors:** Louise L Hardy, Rona MacNiven, Tuguy Esgin, Seema Mihrshahi

## Abstract

**Background:** In New South Wales (Australia) there has been substantial long term investment in school-based child obesity prevention programs. Whether these programs have led to population level improvements in children’s adiposity and weight-related behaviours in Aboriginal children, who are at greater risk of poorer health outcomes, is yet to be determined. The purpose of this study was to describe changes in adiposity and weight-related behaviours of Aboriginal and non-Aboriginal children and to examine the equality of changes between the two groups.

**Methods:** Representative cross-sectional population surveys conducted in 2010 and 2015 among children age 5-16 years (n=15,613), stratified by Aboriginality. Indicators of weight-related behaviour (diet, physical activity, school travel, screen-time) were measured by questionnaire with parents responding for children age <10 years and self-report by children age >10 years. Objective measurements included height, weight, waist circumference, cardiorespiratory fitness, and fundamental movement skills.

**Results:** Adiposity prevalences were significantly higher in 2015, than 2010 among non-Aboriginal children only, however adiposity prevalences were consistently higher among Aboriginal than non-Aboriginal children. There were positive changes towards adopting healthier weight-related behaviours in all children between surveys, which were consistently significant among non-Aboriginal, but not Aboriginal, children. The magnitude of changes and the 2015 prevalences in weight-related behaviours were generally similar for Aboriginal and non-Aboriginal children, however positive changes in fruit consumption and locomotor skills were significantly larger among Aboriginal than non-Aboriginal children. The prevalence of being driven to school in 2015 was significantly higher than 2010 for both groups.

**Conclusions:** Overall, there are signs that Aboriginal and non-Aboriginal children are shifting towards healthier weight-related behaviours. However, many unhealthy weight-related behaviours remain highly prevalent. Our findings may have utility for the direction of future health policy and service delivery to Aboriginal and non-Aboriginal children and the development of health promotion programs to build on these improvements in health behaviours.

## Introduction

Since 2002 there has been substantial investment in New South Wales (NSW) Australia to reduce child obesity through a succession of state plans, policies, and programs to support the healthy development of children from 0-18 years.[1–3] The NSW government strategy was to encourage and support opportunities for the community to be healthy through the delivery of evidence-based, interactive, and relevant programs. Within the education sector, child obesity prevention has been addressed through a steady implementation of specific state level mandatory policies and recommended programmes on healthy eating and physical activity including teacher resources and funding support in early childhood settings[4] and schools.[5–8]

Scaling-up school health programs is challenging, and the realisation of health benefits of these programs can take many years. A recent study showed that there has been an increase in the proportion of schools adopting healthy eating and physical activity programs and policies between 2006 and 2013.[9] An important knowledge gap is whether these programs have achieved their objective to improve children’s healthy eating and physical activity and subsequent reduction in child obesity rates. In contrast to controlled small intervention studies, the rigorous evaluation of scaled-up programs is not feasible. Rather, serial population health surveillance surveys can be used to provide estimates of prevalence, change and trends in health and health outcomes.

Data from the last two NSW representative child health surveys showed that the unadjusted prevalence of overweight/obesity of children age 5-16 years has not significantly changed between 2010 and 2015 (23.2% vs 24.5%, respectively)[10] suggesting the net effect of community and school-based investments may be influencing child obesity rates. However, the epidemiological evidence shows that children living in communities with social and economic disadvantage remain at greater risk of overweight/obesity than children from socio-economic advantage.[11] Secondary sub-population analyses of the NSW child surveys showed there are clear and persistent socio-economic and cultural inequalities in the distribution of overweight/obesity in children, and these disparities have increased over time.[12, 13]

In Australia, Aboriginal people are over-represented amongst disadvantaged groups which has contributed to worse health outcomes (including obesity) and lower life expectancies than non-Aboriginal populations.[14] Recent estimates show that overweight/obesity is higher among Aboriginal than non-Aboriginal children.[15] While many factors contribute to this difference, a key contributor to decreasing the incidence and prevalence of overweight and obesity in Aboriginal adult population is through effective community engagement and using Indigenous ways of working.[16]. This was demonstrated in a study that examined the motivators and barriers to exercise in the Perth Noongar Aboriginal community in Western Australia.[16] There are no data on whether NSW investments into the prevention of child obesity have influenced in changes in weight-related behaviours equally among Aboriginal and non-Aboriginal children. To address this evidence gap, we identify the proportion of children exposed to government healthy eating and physical activity in 2015 survey by Aboriginality. We then describe changes in adiposity and weight-related behaviours of Australian Aboriginal and non-Aboriginal children and examined the equality of changes between the two groups using data population health data collected in 2010 and 2015.

## Methods

The data come from repeat, cross-sectional representative, population monitoring surveys of NSW children age 5-16 years that were conducted between February and March in 2010 and 2015. The survey methodology, including sampling frame, sample size calculations, staff and indicator measurement protocols have remained consistent across survey years to allow comparability and generation of trend information which are described in detail in the survey reports.[10, 17]

Briefly, the sampling frames comprise of primary and secondary schools randomly selected from each education sector (government, Catholic and independent) across socio-economic quintiles and in urban and rural areas, to provide a representative sample of NSW school-age children. The sampling frames were based on a two-stage probability sample (school, student) and comprised all NSW schools, except special education (e.g., blind, sport) and small (<180 enrolments) schools. Schools were randomly sampled from each education sector proportional to enrolment in that sector, then two classes randomly selected from each target year and children in those classes invited to participate. The surveys were school-based, and the data collected by trained teams of teachers. Ethics approvals were given by The University of Sydney, NSW Department of Education and Communities, the NSW Catholic Education Commission and the Aboriginal Health and Medical Research Human Ethics Committee. Written consent from parents was a requirement for participation.

The primary purpose of the surveys are to report on the change in rates of overweight/obesity and weight related behaviours in relation to the NSW State Health Plan target.[3] Cluster sampling was employed for the surveys to adjust for the correlation of measures for children in the same school. This required that the sample size be inflated to account for the clustering effect. The highest cell prevalence for obesity/overweight in 2010 was 29.9% and sample size was calculated using p1=0.30 and p2=0.20, which enabled detection of a difference of 10% in the prevalence between groups, with 80% power and alpha=0.05. Post-stratification weights were calculated to permit inferences from children included in the sample to the populations from which they were drawn, and to have tabulations reflect estimates of population totals. Post-stratification weights were calculated using the NSW student population frame provided by the Australian Council for Educational Research.

## Measures

Adiposity was determined from measured height (m), weight (kg) and waist circumference (cm) using standard procedures.[18] Body mass index (kg/m2) was calculated and children categorised according to international paediatric cut-points [19] as overweight/obese or obese. Abdominal obesity, an index of cardio-metabolic risk, was determined by the waist-to-height ratio (WHtR, cm/cm) and children categorised as low risk (<0.5) or high risk (≥0.5).[20]

Parents of children in Kindergarten, years 2 and 4 (i.e., age <10 years) completed a questionnaire on their child’s demographic variables and indicators of weight-related behaviours distributed at the time of consent. Children in years 6, 8 and 10 (i.e., age >10 years) completed the same questionnaire during the school visits. Demographics included sex, date of birth, postcode of residence, and Aboriginality. Postcode of residence was used as a proxy for socioeconomic status (SES), based on the Australian Bureau of Statistics’ Index of Relative Socio-economic Disadvantage (IRSD). IRSD is one of the socioeconomic indices for areas, which is updated after each Census. The IRSD is an ordinal measure (based on a standard score of 1000 with a standard deviation of 100). For SES we calculated the mean IRDS and quintiles. [21] Postcode of residence was also used to define residence (urban or rural) using the Australian Statistical Geography Standard Volume 5 – Remoteness Areas.[22] 2011 Census data were used for both surveys. Aboriginality was determined by asking *Are you of Aboriginal and*/*or Torres Strait Islander origin?* (*Response; Yes, No, Don’t know*). Don’t know responses were recoded as No (1.9%).

Indicators of dietary intake were collected using a validated short food frequency questionnaire developed for population-based surveys.[23] Respondents reported the usual consumption of fruit, vegetables, energy-dense, nutrient poor foods (EDNP, i.e., fried potato products, salty snack foods, snack foods, confectionary, icecream). The response categories were never/rarely, 1-2 times/week, 3-4 times/week, 5-6 times/week, daily, and 2 or more times/day. For the analysis, a junk food intake measure score was calculated from the combined consumption of EDNP foods and children’s scores ranked into tertiles.[24] Additional dietary behaviours questions included frequency of eating breakfast (daily vs not daily), eating dinner in front of the TV (<5 vs ≥5 times/week), and how often good behaviour was rewarded with sweets (never vs usually/sometimes). Recreational screen-time was measured by questionnaire[25] and children were categorised according to screen-time recommendation (<2 or ≥2hours/day).[26] Respondents also reported if there was a TV in the child’s bedroom (yes/no) and whether there were rules on screen-time (usually vs never/sometimes).

Cardiorespiratory fitness was assessed in children in Year 4 and above using the 20-metre shuttle run test.[27] Children were categorised as ‘adequately fit’ or ‘unfit’ according to the FITNESSGRAM cut-points.[28] Seven fundamental movement skills (FMS) were assessed; four locomotor skills (sprint run, vertical jump, side gallop, leap) and three object control skills (catch, over arm throw, kick) using process-oriented checklists for each skill.[29] Students who demonstrated advanced skills in ≥2 object control and ≥3 locomotor skills were categorized as having advanced object control and locomotor skills, respectively.[30]

In 2015, the Principal of each participating school (primary n=42; secondary n=42) completed a School Environment Question, which included a question on the schools’ participation in established government school-based initiatives to address child obesity in NSW.[10] Responses for each school were linked to children’s data to determine their exposure to these initiatives at school.

## Analysis

We used Stata Version 14.2 (College Station, TX) survey commands to incorporate the sampling weights and to account for the cluster design of the study and adjust for the standard errors and produce 95% confidence intervals. Alpha was set at .05. Missing data were not replaced. Chi-squared tests and tests of independence were used to assess the differences between Aboriginal and non-Aboriginal children’s children exposed to NSW government school-based initiatives. Generalised linear models with a binomial distribution and log link were used to generate prevalences with 95% confidence limits (95%CI) by Aboriginality and survey year, adjusted for sex, SES, residence and age/year. Prevalence ratios (PR, a measure of relative difference) with 95% confidence limits adjusted for sex, SES, residence and age/year and an interaction term between survey years and Aboriginality status were generated to quantify the differences between survey years for Aboriginal and non-Aboriginal children. We also used PR (95%CI) to estimate differences in the change in prevalence between surveys between Aboriginal (reference group) and non-Aboriginal children, adjusted for sex, SES, residence and age, and an interaction term between survey year and Aboriginality status. The interaction term represents the effect of time on change in weight and weight-related behaviours for Aboriginal children relative to non-Aboriginal children.

## Results

The characteristics of the children are presented in Table 1 by Aboriginality and survey year. The survey response rates were 57% and 62% in 2010 and 2015, respectively. Aboriginal children comprised 3.2% and 3.4% of the sample in 2010 and 2015, respectively. Aboriginal children were more likely to live in rural areas and have lower SEIFA scores than their non-Aboriginal peers in both survey years. Table 2 shows the number of schools with different healthy eating and physical activity programs and the overall proportion of Aboriginal and non-Aboriginal children’s who have exposure to these programs. There were no statistical differences between Aboriginal and non-Aboriginal children’s exposure to these programs, however there were differences in program adoption by schools. In primary schools the most prevalent were fruit/vegetable/water programs and in high schools the inclusion of healthy eating and physical activity in the curriculum.

**Table 1.** Sample characteristics by Aboriginality and survey year.

|  | Aboriginal |  |  | Non-Aboriginal |  |  |
| --- | --- | --- | --- | --- | --- | --- |
| Survey year | 2010 | 2015 | P-value | 2010 | 2015 | P-value |
| n (%)* | 254 (3.2%) | 251 (3.4%) |  | 7,594 (96.8%) | 7,191 (96.6%) | .65 |
| Girls (%; 95%CI) | 55.8 (51.5, 59.9) | 46.8 (40.3, 59.7) | .03 | 47.7 (46.6, 48.9) | 50.6 (47.1, 54.1) | .12 |
| Age (years, range) | 10.7 (11.6) | 10.3 (12.2) | .15 | 10.6 (4.2,) | 10.3 (13.9) | <.001 |
| SES (mean, range)** | 941 (565) | 983 (375) | <.001 | 989 (573) | 1000 (576) | <.001 |
| *Residence (%; 95%CI)* |  |  |  |  |  |  |
| Urban | 61.3 (59.9, 62.6) | 65.1 (48.6, 78.6) | .63 | 85.1 (82.0, 87.7,) | 76.7 (65.9, 84.8) | .06 |
| Rural | 38.7 (37.4, 40.1) | 34.9 (21.4, 51.4) |  | 14.9 (12.3, 18.0) | 23.3 (15.2, 34.1) |  |

\* Aboriginal children; 2010 survey primary school n=157, secondary school n=97; 2015 survey primary school n=176, secondary school n=75.
\*\* SES = Socioeconomic status. Based on the Australian Bureau of Statistics' Index of Relative Socio-economic Disadvantage

**Table 2.** Proportion of children attending schools in 2015 with government school-based child obesity prevention initiatives, (reported by the Principal) by school level and Aboriginality*

| NSW government school-based child obesity initiatives | Schools<br>participating<br>(n) | Children |  |
| --- | --- | --- | --- |
|  |  | Aboriginal<br>(%, 95%CI) | Non-Aboriginal<br>(%, 95%CI) |
| *Primary school (n)* | 44 | 176 | 4,969 |
| Fruit, vegetable, or water breaks | 41 | 96.4 (85.3, 99.2) | 94.8 (80.7, 98.8) |
| Crunch and Sip®** | 27 | 63.5 (39.0, 82.6) | 60.4 (44.4, 74.5) |
| School kitchen garden | 9 | 34.5 (16.7, 57.9) | 43.3 (28.0, 59.9) |
| Q4: H2O (a student activity to promote healthy drinks)* | 9 | 20.7 (7.7, 45.0) | 20.1 (10.0, 36.1) |
| Heart Foundation Jump Rope for Heart® | 15 | 33.9 (10.7, 16.2) | 31.9 (7.5, 18.9) |
| NSW Premier's Sporting Challenge | 22 | 45.9 (26.2, 66.9) | 51.5 (39.6, 63.3) |
| Live Outside the Box (Student activity to raise awareness about healthy eating, physical activity, limiting screen-time)* | 6 | 17.9 (5.9, 43.4) | 14.5 (6.4, 29.8) |
| Live Life Well @ School initiatives (child obesity prevention program) | 27 | 53.9 (30.1, 76.0) | 57.8 (41.5, 72.6) |
|  |  | Aboriginal<br>(%, 95%CI) | Non-Aboriginal<br>(%, 95%CI) |
| Active After Schools Community | 18 | 33.6 (16.6, 56.2) | 38.0 (24.2, 54.2) |
| Outside of School Hours (OOSH) programs | 27 | 51.2 (28.2, 73.7) | 63.7 (47.3, 77.5) |
| *High school initiatives* | 45 |  |  |
| Children (n) |  | 75 | 2,222 |
| Communication to parents regarding the importance of physical activity in children (usually) | 11 | 21.1 (9.2, 41.4) | 27.1 (15.4, 43.1) |
| Physical activity included in PDHPE scope and sequence (usually) | 42 | 97.7 (84.2, 99.7) | 97.8 (85.1, 99.7) |
| School Plan includes promoting physical activity to students (usually) | 16 | 28.7 (14.1, 49.8) | 37.6 (23.7, 53.8) |
| Leadership programs for older students to encourage physical activity with younger students (usually) | 8 | 17.2 (6.3, 38.9) | 16.8 (8.2, 31.3) |
| Healthy eating included in PDHPE scope and sequence (yes) | 42 | 99.1 (92.9, 99.9) | 97.7 (86.1, 99.7) |
|  |  | Aboriginal<br>(%, 95%CI) | Non-Aboriginal<br>(%, 95%CI) |
| Address healthy eating in the School Plan (yes) | 12 | 43.7 (21.1, 69.2) | 45.4 (28.9, 63.1) |
| Have healthy fundraising options (yes) | 13 | 26.8 (11.0, 51.9) | 40.1 (24.8, 57.6) |
| Encourage the provision of healthy food/drinks to students for activities (e.g., excursions, camps, sport carnivals) (yes) | 27 | 75.1 (53.3, 88.8) | 80.2 (63.1, 90.6) |
| Promote and model healthy eating during school activities involving the wider community, including school fetes and school functions (yes) | 17 | 40.9 (21.0, 64.3) | 49.7 (33.3, 66.2) |
| Communicate to parents the importance of healthy eating (yes) | 14 | 51.2 (27.4, 74.5) | 46.9 (29.4, 65.2) |

\* All $P > .05$ (no difference between groups); \*\*Crunch and Sip® is a registered fruit and water program for schools which provides information and resources including materials for children to take home to their families.
\* These programs were disseminated in only two local health districts, not state-wide.

The change in weight and weight-related behaviours between 2010 and 2015 by Aboriginality are shown in Table 3. Missing data overall was 2.6% for adiposity outcomes, 2.9-3.4% for dietary outcomes, 1.1-5.2% for physical activity outcomes, and 3.2-3.9% for screen-time outcomes. Adiposity prevalences were higher in 2015 than 2010, significantly so for non-Aboriginal children and were higher among Aboriginal than non-Aboriginal children in 2010 and in 2015. Changes in the prevalence of weight-related behaviours were generally significant for non-Aboriginal and not significant for Aboriginal children. However, the overall the pattern of the direction of the change prevalence of many indicators were similar for Aboriginal and non-Aboriginal children. Less healthy changes in all children included a higher prevalence in 2015 of being driven to school, and lower prevalences for daily breakfast, adequate fitness, and active school travel, than in 2010. Healthy changes in all children included a higher prevalence in 2015 of meeting fruit and vegetable recommendations, eating dinner in front of the TV less than 5 times/week and not having a TV in the bedroom, than in 2010.

**Table 3.** Adjusted prevalence of weight and weight-related behaviours by Aboriginality and survey year (%; 95%CI)

|  | **Aboriginal children** |  | **Non-Aboriginal children** |  |
| --- | --- | --- | --- | --- |
|  | 2010<br>(n=254) | 2015<br>(n=251) | 2010<br>(n=7,594) | 2015<br>(n=7,191) |
| *Weight status* |  |  |  |  |
| Overweight/obese | 25.8 (19.8, 31.7) | 29.3 (23.7, 34.9) | 22.1 (20.6, 23.6) | **25.3 (23.7, 27.0)** |
| Obese | 8.0 (4.2, 12.0) | 9.8 (5.8, 13.8) | 5.5 (4.7, 6.2) | **6.9 (6.0, 7.8)** |
| Waist-to-height ratio $\geq 0.5$ | 15.2 (10.5, 20.0) | 17.9 (13.18, 22.8) | 10.1 (8.8, 11.4) | **14.3 (12.6, 16.0)** |
| *Food consumption* |  |  |  |  |
| Mets fruit recommendation | 63.8 (57.0, 70.7) | **76.0 (70.3, 81.6)** | 74.9 (72.9, 76.8) | **78.4 (77.0, 82.0)** |
| Mets vegetable recommendation | 4.2 (1.8, 6.8) | 7.3 (4.2, 10.5) | 4.9 (4.2, 5.6) | **6.8 (6.0, 7.6)** |
| Eat breakfast daily | 73.2 (67.1, 79.3) | 69.7 (64.7, 74.7) | 78.8 (77.3, 80.3) | 76.8 (75.2, 78.5) |
| Junk Food Index Measure |  |  |  |  |
| Low (score 0-5) | 39.7 (33.7, 45.6) | 40.4 (34.5, 46.2) | 39.4 (38.3, 40.5) | **42.9 (41.8, 43.9)** |
| Middle (score 6-8) | 34.3 (28.8, 40.0) | 29.8 (24.7, 35.0) | 29.4 (28.3, 30.5) | 29.0 (28.0, 30.1) |
| High (score 9-25) | 19.4 (14.1, 24.7) | 24.0 (19.6, 28.3) | 23.3 (22.4, 24.2) | 21.8 (20.8, 22.9) |
| *Family practices* |  |  |  |  |
| Eats dinner in front of TV <5/week | 75.0 (68.6, 81.5) | 81.1 (76.6, 85.5) | 80.2 (78.8, 81.5) | **82.6 (81.5, 83.8)** |

|  | Aboriginal children |  | Non-Aboriginal children |  |
| --- | --- | --- | --- | --- |
|  | 2010<br>(n=254) | 2015<br>(n=251) | 2010<br>(n=7,594) | 2015<br>(n=7,191) |
| Parents never reward good behaviour with sweets | 41.4 (35.3, 47.5) | 44.3 (37.6, 50.9) | 48.8 (47.0, 50.5) | **46.2 (44.4, 47.9)** |
| *Physical activity outcomes* |  |  |  |  |
| Advanced locomotor skills | 27.5 (21.3, 33.7) | **48.8 (42.0, 55.6)** | 35.7 (33.5, 38.0) | **48.0 (45.5, 50.5)** |
| Advanced object control skills | 46.0 (41.7, 50.4) | 47.0 (43.3, 50.1) | 42.8 (40.9, 44.8) | **45.1 (43.1, 47.1)** |
| Healthy cardiorespiratory fitness zone | 66.5 (58.9, 74.0) | 56.4 (48.8, 64.1) | 66.2 (63.0, 69.4) | **60.5 (56.8, 64.2)** |
| *School travel* |  |  |  |  |
| Active travel to school | 20.3 (14.4, 26.2) | 14.2 (8.4, 20.0) | 16.2 (13.8, 18.7) | 13.7 (11.6, 15.9) |
| Driven to school | 23.8 (16.1, 31.5) | **38.2 (30.5, 45.8)** | 36.1 (32.7, 39.5) | **44.9 (41.6, 48.3)** |
| *Screen time* |  |  |  |  |
| Met recommendation on weekdays | 39.9 (33.4, 46.4) | 46.9 (40.8, 53.1) | 54.5 (52.4, 56.5) | 52.5 (49.9, 55.1) |
| Met recommendation on weekends | 18.5 (12.5, 24.5) | 20.0 (14.3, 25.6) | 17.7 (16.2, 19.1) | 19.5 (17.9, 21.1) |
| No TV in the bedroom | 47.5 (39.6, 55.3) | 56.2 (49.9, 62.5) | 72.3 (69.9, 74.6) | **77.9 (75.8, 80.0)** |
| Usually has rules on screen-time | 40.1 (32.1, 48.1) | 47.6 (40.8, 54.5) | 49.3 (47.5, 51.0) | **46.4 (44.8, 48.0)** |

Bold values represent significant difference between 2010 and 2015 within each child group. Models adjusted for sex, age, residence (rural/urban), and socioeconomic quintile.

Table 4 shows the relative change in weight and weight-related behaviours between surveys by Aboriginal status and for Aboriginal compared with non-Aboriginal children. Between 2010 and 2015, there were increases in adiposity, but these were only statistically significant for non-Aboriginal children, where overweight/obesity increased by 15%, obesity by 25% and WtHR by 42%. Overall, there were seven significant positive improvements in weight related behaviours of non-Aboriginal children and two significant positive changes in Aboriginal children. Aboriginal children’s fruit intake was 14% higher than the increase in non-Aboriginal children, but only marginally (95%CI 1.00, 1.29) and the increase in advanced locomotor skills was 32% (95%CI 1.02, 1.70) higher than non-Aboriginal children. Less healthy behaviour changes between 2010 and 2015 included a 63% (95%CI 1.11, 2.38) and 24% (95%CI 1.09, 1.40) increase in being driven to school, among Aboriginal and non-Aboriginal children, respectively. Among non-Aboriginal children cardiorespiratory fitness and usually having rules on ST decreased 9% and 6%, respectively, between survey years. Although physical activity data were not available for 2010, in 2015 26.8% of Aboriginal children met the recommendation for daily physical activity, compared with 18.8% of non-Aboriginal children (PR 1.58, 95%CI 1.05 to 2.35) (data not shown).

**Table 4.** Prevalence ratios (PR) of change in weight and weight-related behaviours for Aboriginal and non-Aboriginal children by survey year and Aboriginality (PR 95%CI)

|  | Prevalence ratio (95%CI) |  |  |
| --- | --- | --- | --- |
|  | Aboriginal children | Non-Aboriginal children | Aboriginal (ref)/non-Aboriginal<sup>2</sup> |
| Indicator of behaviour | 2010/2015<sup>1</sup> | 2010/2015<sup>1</sup> |  |
| *Weight status* |  |  |  |
| Overweight/obese | 1.14 (0.84, 1.54) | **1.15 (1.04, 1.26)** | 0.99 (0.74, 1.33) |
| Obesity | 1.24 (0.67, 2.30) | **1.25 (1.04, 1.52)** | 0.99 (0.53, 1.83) |
| Waist-to-height ratio $\geq 0.5$ | 1.18 (0.77, 1.80) | **1.42 (1.19, 1.71)** | 0.83 (0.55, 1.25) |
| *Dietary intake* |  |  |  |
| Mets fruit recommendation | **1.19 (1.04, 1.36)** | **1.05 (1.02, 1.08)** | 1.14 (1.00, 1.29) |
| Mets vegetable recommendation | 1.74 (0.84, 3.60) | **1.40 (1.17, 1.67)** | 1.25 (0.61, 2.56) |
| Eats breakfast daily | 0.95 (0.85, 1.06) | 0.97 (0.95, 1.00) | 0.98 (0.88, 1.09) |
| *Junk Food Index Measure* |  |  |  |
| Low | 1.02 (0.83, 1.25) | **1.09 (1.05, 1.13)** | 0.93 (0.76, 1.16) |
| Middle | 0.87 (0.69, 1.10) | 0.99 (0.94, 1.04) | 0.88 (0.69, 1.12) |
| High | 1.23 (0.89, 1.72) | 0.94 (0.88, 1.00) | 1.32 (0.94, 1.85) |
| *Family practices* |  |  |  |
| Eats dinner in front of TV <5 times/week | 1.08 (0.98, 1.20) | **1.03 (1.01, 1.05)** | 1.05 (0.95, 1.16) |
|  | Aboriginal children | Non-Aboriginal children | Aboriginal (ref)/non-Aboriginal<sup>2</sup> |
| Indicator of behaviour | 2010/2015<sup>1</sup> | 2010/2015<sup>1</sup> |  |
| Parent's never reward good behaviour with sweets | 1.07 (0.86, 1.32) | 0.95 (0.90, 1.00) | 1.13 (0.92, 1.39) |
| *Physical activity outcomes* |  |  |  |
| Advanced locomotor skills | **1.77 (1.36, 2.31)** | **1.34 (1.24, 1.45)** | **1.32 (1.02, 1.70)** |
| Advanced object control skills | 0.98 (0.86, 1.12) | **1.75 (1.01, 1.13)** | 0.92 (0.80, 1.05) |
| Healthy cardiorespiratory fitness zone | 0.85 (0.71, 1.01) | **0.91 (0.85, 0.99)** | 0.93 (0.78, 1.11) |
| *School travel* |  |  |  |
| Active travel to school | 0.70 (0.42, 1.16) | 0.85 (0.67, 1.07) | 0.82 (0.50, 1.37) |
| Driven to school | **1.63 (1.11, 2.38)** | **1.24 (1.09, 1.40)** | 1.31 (0.91, 1.89) |
| *Screen time* |  |  |  |
| <2hrs ST weekdays | 1.17 (0.95, 1.44) | 0.96 (0.91, 1.02) | 1.21 (1.00, 1.49) |
| <2hrs ST weekends | 1.08 (0.70, 1.67) | 1.10 (0.98, 1.24) | 0.98 (0.63, 1.51) |
| No TV in the bedroom | 1.18 (0.97, 1.45) | **1.08 (1.03, 1.13)** | 1.10 (0.90, 1.34) |
| Usually has rules on ST | 1.19 (0.93, 1.51) | **0.94 (0.90, 0.99)** | 1.26 (0.99, 1.61) |

\*Adjusted for sex, age/year group, socioeconomic quintile, residence (urban/rural); bold values = significant; <sup>1</sup> Reference group = 2010; <sup>2</sup> Reference group = Aboriginal children

## Discussion

The purpose of this study was to explore population changes in children’s adiposity and weight-related behaviours following more than a decade of sustained, evidence-based, scaled-up child obesity prevention programs, and to examine these changes among Australian Aboriginal children compared with non-Aboriginal children. Our findings indicated adiposity has increased among NSW children between 2010 and 2015, however there were positive changes in many weight-related behaviours, the precursors for change in adiposity. While the changes in prevalences were generally significant for only non-Aboriginal children, the pattern and magnitude of change in prevalences were similar among Aboriginal children. The lack of statistical significance needs to be considered in the context of the small sample of Aboriginal children which was a limitation of the surveys and potentially under powered our analyses as evident by the wide confidence intervals. Although the survey sampling frame was designed to represent NSW children age 5-16, not Aboriginal children, Aboriginal children represented 3.5% of the sample survey which approximates the state population prevalence of 5.8% in NSW.[31] Another limitation was the use of self-report, but the questionnaires used validated indicators of weight-related behaviours.

Despite these limitations the findings have relevance to the health of Aboriginal children. Addressing the health inequality gap between Aboriginal and non-Aboriginal people is an Australian government priority[32] and population health surveys give an indication of whether there is progress towards addressing health inequities. An encouraging key finding was that there was no difference between Aboriginal and non-Aboriginal children’s exposure to school-based child obesity prevention programs on healthy eating and physical activity. However, except for fruit/vegetable/water programs in primary schools, the adoption of many government child obesity prevention programs in schools was low. This is consistent with other studies that suggest additional and/or different dissemination strategies are required to facilitate a greater adoption of policies and practice within schools.[9, 33]

While we found there were positive changes in many weight-related behaviours which were generally in the same direction and of the same magnitude for Aboriginal and non-Aboriginal children, adiposity prevalences were higher in 2015 than 2010 for all children and were consistently higher among Aboriginal children. This may indicate there are issues with the fidelity and quality of the delivery of these programs in some schools. Alternatively, school-based programs alone may not change weight-related behaviours in the absence of efforts to change the broader obesogenic environment, which may have greater influence in disadvantaged communities.[34]

Two notable improvements among Aboriginal compared with non-Aboriginal children were fruit intake and locomotor skills. School-based initiatives to improve children’s fruit/vegetable intake and FMS have been scaled up across NSW early childhood settings and primary schools for more than a decade. It is therefore reasonable to conclude that schools have been a major contributor to the temporal improvements in these behaviours, particularly for Aboriginal children. In 2005, in response to low fruit and vegetable consumption, the Australian Government recommended all primary schools implement a vegetable and fruit break program.[35] In this study >95% of primary school children were exposed to these programs and the 14% increase in Aboriginal children’s fruit intake suggests these programs may have played an important role in this increase.

Similarly, since 2008, in response to children’s low proficiency in FMS, government and independent groups have delivered FMS programs across early childhood settings in NSW.[7] The prevalence of primary school children’s FMS proficiency has significant increased between 2010 and 2015,[10] suggesting FMS programs during preschool and primary school years have been beneficial to children’s acquisition of FMS. While not all children attend formal early childcare, the number of Aboriginal children attending formal preschool has been increasing,[36] thus increasing the likelihood of their child’s exposure to a FMS program.

The relationship between FMS and physical activity is reciprocal [37] which may help explain the increase in Aboriginal children’s locomotor skills. Our findings are consistent with national data which show Aboriginal children are generally more active than non-Aboriginal children.[38] Aboriginal community workers have suggested possible factors for their children’s higher activity include more children in the community family unit to play with and that play is valued and encouraged.[39] The lower prevalences of cardiorespiratory fitness, in both child groups, in 2015 compared with 2010 are however concerning and may indicate the need to target future large scale programs, policies and promotional efforts. We were unable to examine change in children’s physical activity because the question changed between surveys, however in 2015 children’s physical activity levels were low with few meeting the daily 60-minute recommendation. This may have contributed to the decline in cardiorespiratory fitness between surveys.

Our findings also highlighted several behaviours that require further investment to improve. Although not statistically significant, there was a decrease in daily breakfast consumption among both groups of children. The association between skipping breakfast and adiposity is contentious,[40] however there is evidence that children who skip breakfast tend to have poorer diet quality.[41] Our findings showed an increase between survey years in the prevalence of a high junk food intake, but only among Aboriginal children. While we did not measure food insecurity, it has been identified as a leading health issue in Aboriginal communities and is associated with consuming foods of poor quality because they are the lowest cost options.[42] A recent review identified that effective nutrition programs involved strong Aboriginal community engagement in the development and implementation of programs to community priorities are addressed.[43] The implementation of such programs in schools with high Aboriginal student populations could have merit, but other social and cultural determinants such as income must also be addressed to improve Aboriginal children’s dietary intakes.

There were significant increases in the prevalence of being driven to school among both child groups, which potentially reduces children’s overall daily physical activity.[44] Reasons for the increase in being driven to school are not clear, but other Australian research suggests factors associated with children not using active transport to/from school include proximity to school, the safety of the route, and family time constraints.[45] Recent Australian studies have reported that many children did not attend the school closest to them because their parents preferred to send them elsewhere, and that increase in chauffeuring children, particularly to school has led to declines in children’s independent mobility.[46, 47]

Schools play an important role in health promotion, however the origins of many weight-related behaviours begin in the home, and for meaningful population behaviour change in children interventions that involve parents are necessary, particularly among disadvantage families. In NSW obesity prevention initiatives with a focus on Aboriginal children are limited, but there are an increasing number of programs include Aboriginal content.[48, 49] Unfortunately while many programs collect evaluation data, publication of findings are underrepresented in academic literature.[48] A lack of evidence may potentially hamper health promotion efforts in Aboriginal communities.

The strengths of this study include the consistent sampling frames and protocols which allowed comparison across survey periods and the generalisability of findings to children age 5-16 years living in NSW. Further, this is the first study to examine whether indicators of adiposity and weight-related behaviours have changed equally across NSW Aboriginal and non-Aboriginal children over time, with measured anthropometry, and validated indicators of weight-related behaviours.

## Conclusions

Community-wide programs are required to shift population behaviour, and it is important to remember they take time to implement across a range of different population groups, settings, and sectors, and even longer to yield results. Overall, this study found that there are signs that Aboriginal children are shifting towards healthier weight-related behaviours. Many unhealthy weight-related behaviours remain highly prevalent and reducing adiposity prevalence in all children is contingent upon sustained investment in preventive health initiatives to enable the adoption of healthier weight-related behaviours. Our findings may have utility for the direction of future health policy and service delivery to Aboriginal children in particular and the development of health promotion programs to build on these improvements in health behaviours.

## Acknowledgements

We thank the schools and students for their participation.

